# A Novel Therapeutic Approach: Gamma Secretase Inhibitor Enhances Radiotherapy and Checkpoint Blockade Therapy via Reprogramming of the Tumor Microenvironment

**DOI:** 10.64898/2025.12.30.696685

**Authors:** Qi Wang, Debarshi Banerjee, Erin C. Connolly, Claire Vanpouille-Box, Carrie Shawber, Rami Vanguri, Michael Kissner, Michael L. Miller, Darrell J Yamashiro, Eileen P. Connolly

## Abstract

**Background:** High-dose radiotherapy (RT) in cancer is immunogenic but also induces an immunosuppressive tumor microenvironment (TME) that limits the efficacy of immune checkpoint inhibitors (ICIs). Overcoming this immunosuppressive barrier is therefore critical to unlocking the full potential of RT-ICI combinations. As Notch signaling regulates tumor microenvironment, we hypothesized that the γ-secretase inhibitor (GSI), AL101, would suppress radiation-induced immunosuppression and enhance antitumor efficacy when combined with RT and anti-PD-1 (aPD-1) therapy.

**Methods:** Syngeneic neuroblastoma (9464D) and triple-negative breast cancer (EO771) tumors were established in C57BL/6, macrophage-depleted C57BL/6 mice, or athymic mice. Mice received 12 Gy RT (day 3), AL101 (6.5 mg/kg daily, day 0-9), and aPD-1 (days 0, 3, 6). Tumors were analyzed by spectral flow cytometry and single-cell RNA sequencing (scRNA-seq), and lung metastases were evaluated histologically.

**Results:** The triple combination of RT, aPD-1, and GSI produced durable tumor growth inhibition and significantly prolonged survival in both models, with median survival more than doubled compared to all other treatment groups. Triple therapy also markedly reduced lung metastases in EO771 mice. These effects were abrogated in athymic nude mice and macrophage-depleted immunocompetent mice, consistent with an immune-dependent mechanism. Based on scRNA-seq and spectral flow cytometry analysis, RT alone increased exhausted T cells and immunosuppressive macrophages, while triple therapy reversed these effects, including expansion of activated CD8⁺ T cells, reduction of Tregs and exhausted T cells, restoration of cross-presenting CD103⁺ dendritic cells, and reprogramming of myeloid cells toward a proinflammatory state.

**Conclusions:** GSI remodels the RT-induced immunosuppressive TME and potentiates the efficacy of RT + ICI. GSI combined with RT+aPD-1 reprograms the TME toward an immunostimulatory state and supports GSI as a promising immuno-radiotherapeutic strategy with strong translational potential.

## Introduction

Immune checkpoint inhibitors (ICIs) targeting PD-1 or PD-L1 activate the immune system to reject tumors, but not all patients respond ^1, 2^. Although RT can be immunogenic and enhance ICI efficacy preclinically^3, 4^, this treatment also recruits immunosuppressive cells into the tumor microenvironment (TME), including Tregs^5^ and tumor-associated macrophages (TAMs)^6, 7^ that limit response durability^8^. Strategies that amplify RT-induced immunogenicity while countering radiation-driven immunosuppression have been proposed to improve outcome with RT + ICI.

The Notch family consists of four transmembrane receptors (Notch1-4) and five transmembrane ligands (Dll1/3/4, Jag1/2)^9, 10^. Notch signaling regulates diverse aspects of cancer biology, including cancer cell metabolic programming, stromal remodeling, and immune cell differentiation^11^. In the TME, its roles are context-dependent: Notch supports anti-tumor immunity by promoting CD8⁺ T cell differentiation and memory^12–14^, dendritic cell maturation^14^ in a cell-autonomous manner, and limiting immunosuppressive TAM recruitment in non-cell-autonomous manner^15, 16^. Conversely, Jag1/Notch signaling expands Tregs^17^ and can skew macrophages toward pro-tumor states^18, 19^, while Notch upregulation of PD-1 promotes T cell exhaustion in CD8⁺ T cells^14, 20^. This duality highlights the complexity of targeting Notch in the TME immune cells^14^.

γ-secretase inhibitors (GSIs) are small-molecules that block cleavage of Notch receptors, thereby preventing release of the Notch intracellular domain and suppressing downstream canonical Notch signaling^21^. Previously we have reported the Notch signaling components of the tumor vasculature upregulated upon the RT in neuroblastoma, a common extracranial pediatric solid tumor^22^. Here, we demonstrated that GSI significantly enhanced the efficacy and reduced metastasis of RT with aPD-1 immunotherapy in two immunologically distinct tumor models (neuroblastoma and breast cancer). We further found that triple therapy reprogramming the tumor immune microenvironment (TIME) through myeloid reprogramming enhanced antigen presentation, and cytotoxic T-cell activation. These findings support GSI as a strategy to overcome RT-induced immunosuppression and potentiate RT + ICI therapy, achieving both primary tumor suppression and metastatic burden reduction.

## Methods

### Cells

EO771 cells (purchased from ATCC) and the 9464D cells (derived from a TH-MYCN transgenic neuroblastoma mouse, obtained as a gift from Dr. Crystal Mackall at the National Institute of Health) were cultured in DMEM with 10% fetal bovine serum (FBS, Euroclone) and 1% non-essential amino acids supplemented with 10% FBS, at 37°C and 5% CO_2_. Both cell lines were authenticated by short tandem repeat profiling (ATCC). Cell derivatives were maintained in DMEM supplemented with10%FBS, at 37°C in 5% CO_2_ and routinely monitored for signs of contamination by visual inspection under phase-contrast microscopy. Cells were used at 80% confluency for tumor injection.

### Clonogenic assay

The clonogenic assay was performed to evaluate the radiation sensitivity of EO771 and 9464D tumor cell lines as previously describe^23^. Briefly, cells were seeded (ranging from 100 cells to 20,000 cells per well, in triplicates) in 6-well plates. Cells were treated with or without GSI (AL101 100nM, 55.65 ng/ml) for 24 h after plating. Then, cells were exposed to increasing doses of ionizing radiation (0, 2, 4, 8, or 12 Gy) using a Gamma-40 irradiator (dose rate 209.3cGy/min). Following irradiation, cells were incubated under standard culture conditions (37°C, 5% CO₂) for 10-14 days to allow colony formation. The resulting colonies were then fixed with methanol and stained with crystal violet. The surviving fraction at each radiation dose was calculated relative to the non-irradiated control (0 Gy) to compare the radiosensitivity of EO771 and 9464D cells.

### Animal model

All procedures performed were approved by the Institutional Animal Care and Use Committee (IACUC) of Columbia University (AABQ7586). 9464D cells (10^6^) were injected subcutaneous into the flanks of female C57BL/6 and NCr nude mice (Taconic). EO771 cells (10^5^) were injected into the mammary fat pad of syngeneic 16-40-week-old female C57BL/6 mice. Tumor growth was measured twice weekly using digital calipers, and tumor volume was calculated as (length × width²)/2. When tumors reached a volume of approximately 150-200mm^3^, mice were enrolled as day 0 of experiment and randomized into treatment groups (5-9 mice/group). For the RT-alone or combination treatment groups, primary tumors were irradiated with a single 12 Gy high-dose fraction on day 3 after enrollment using the Small Animal Radiation Research Platform (SARRP; Xstrahl) ^24^. Image-guided irradiation was performed to precisely target the tumor while shielding the remainder of the body. X-ray irradiation was delivered at 220 kV with a dose rate of 310 cGy/min. CT scans were obtained in four representative mice to define tumor location and guide treatment planning, and the resulting setup was applied uniformly to all remaining animals to ensure consistent dose delivery. Mice received aPD-1 monoclonal antibody (Invivo Mab, 10mg/kg body weight, i.p.) on days 0, 3, and 6 after irradiation, and GSI AL101 (6.5 mg/kg, oral gavage) was administered daily from day 0 through day 9 post-irradiation. Control mice received either vehicle or an isotype antibody (anti-IgG, aIgG) on the same schedule. For macrophage depletion experiments, mice were treated with clodronate liposomes (SUV PEG liposome formulation; Liposoma, The Netherlands; 100 μL per mouse, i.p.), starting 2 days before tumor enrollment. Injections were administered every 5-7 days, beginning prior to treatment initiation and continued throughout the study. Control animals received PBS liposomes on the same schedule. Mice were euthanized when tumor volume exceeded 1500 mm³ or if animals met humane endpoint criteria.

### Flow cytometry

To analyze immune cell subsets, mouse tumors were dissected and minced, then the suspension was filtered. Viability was assessed by the Zombie NIR Dead Cell Stain Kit (Invitrogen Life Technologies). Samples were washed then followed by staining with a master mix of surface antibodies (Supplementary Table 1-2) and incubation. Cells were then fixed using FoxP3 Fix/Perm kit fixation buffer (Thermo Fisher-eBioscience) and incubated on ice. Afterwards, cells were washed with 1X Perm Wash buffer (Thermo Fisher-eBioscience) and then stained with master mix of intracellular antibodies (Supplementary Table 1-2). The cells were resuspended and measured on an ID7000 Spectral Cell Analyzer (Sony Biotechnology). Flow cytometry data were analyzed using FlowJo (v10.7.1). The gating strategy for different cell subpopulations is shown in **Supplementary Fig. 1**.

### Single-cell RNA sequencing

Mouse tumors were dissected, minced, and enzymatically dissociated using DNase and collagenase into single-cell suspensions. The resulting suspensions were filtered through a 70-µm cell strainer, washed with phosphate-buffered saline (PBS), and adjusted to a final concentration of 1-2 × 10⁶ cells per 100 µL. To enrich immune populations, staining was performed using PE/Cyanine7 anti-mouse CD45 antibody (Thermo Fisher). Cells were hashtagged using the CellPlex kit (10x Genomics) and sorted on a SONY MA900 cell sorter (Sony Biotechnology) to isolate viable, CD45⁺ cells. Cells derived from animals within the same experimental group (n=4-5 mice/group) were pooled prior to Gel Beads-in-emulsion (GEM) generation and library preparation following the manufacturer’s protocol. Sequencing data were processed using the Cell Ranger Single Cell Software Suite (v5.0.1, 10x Genomics) for demultiplexing, barcode processing, and 3′ gene counting, with alignment to the mm10 (2020) mouse genome. Count matrices were analyzed in R (v4.2.1) using the Seurat package (v4.1.1). Cells with <500 detected genes, >10% mitochondrial transcripts, >40,000 UMIs, or >5000 detected genes were excluded. Data were log-normalized (NormalizeData, scale factor = 10,000), and the 2,000 most variable genes were identified. Gene expression matrices were scaled using ScaleData after regressing out cell cycle and mitochondrial effects.

Dimensionality reduction was performed using principal component analysis (20 PCs) followed by UMAP visualization. Cell clusters were identified using the Leiden algorithm and annotated based on canonical marker genes. Differential expression analysis was conducted using Seurat’s FindAllMarkers function (Wilcoxon test, min.pct = 0.1, logfc.threshold = 0.5).

To compare cell subset proportions across treatment conditions, we used the scProportionTest package with permutation-based testing. Gene signature enrichment was calculated using the UCell package, and density visualization was generated with the Nebulosa package to illustrate the distribution of signature scores across cell populations.

We quantified M-MDSC transcriptional activity at single-cell resolution by computing a module score from a curated gene set of canonical M-MDSC markers (Arg1, Ly6c, S100a8, S100a9, Il1b, Ptgs2, Itgam) using UCell, a rank-based method robust to sequencing depth variation. Scores were computed on the log-normalized expression matrix and used for visualization and group comparisons across myeloid clusters.

### HALO-AI Lung metastasis analysis

Lung tissues from mice were harvested, fixed in formalin, paraffin-embedded, sectioned at 5 µm, and H&E stained. Whole-slides were scanned using Leica AT2 whole slide digital imaging system for 40x followed by Indica HALO-AI software (version 3.5) MiniNet classifier analysis. The algorithm was configured and validated to accurately detect and quantify metastatic nodules, normal lung tissue and adjacent tissues. Batch processing was performed for consistent analysis across samples.

### Statistics

Survival curves were built by the Kaplan-Meier method and compared using Log-rank (Mantel-Cox) test. Comparison between different groups used one-way ANOVA, followed by Tukey post-hoc comparisons. All experiments including the quantitative data of metastasis were analyzed using Graphpad Prism 10. A p-value <0.05 will be considered significant.

## Results

### Combination RT + aPD-1+GSI significantly improves survival in two distinct tumor models

9464D and EO771 tumor cells were either embedded in the mammary fat pads of C57BL6 female mice or subcutaneously in nude mice, respectively. After tumors reach 100-150mm^3^, mice were divided into groups followed by irradiation and aPD-1 and A101 (GSI) administration as described in Fig 1A. Untreated mice median survival was 12 days in 9464D mice and 14 days in EO771 mice, GSI or aPD-1 monotherapy failed to significantly affect tumor growth or survival (median survival: 9464D GSI:10d, aPD1:10d; EO771 GSI:10d, aPD1:15d), and consist with previous publication^25^, RT produced an intermediate the survival benefit (median survival: 9464D:37d; EO771:35d). Dual-administration of GSI or aPD1 with RT regimens did not substantially improve outcomes beyond RT alone (median survival: 9464D RT+GSI:50d, RT+aPD-1: 46d; EO771:RT+GSI: 41d, RT+aPD-1: 43d) (**Fig. 1B, C**). RT + aPD-1 + GSI triple therapy produced a dramatic survival benefit for more than double of the dual therapy in both tumor models. When compared with RT + aPD-1, triple therapy median survival increase to 87 days versus 46 days in the 9464D model and 98.5 days versus 43 days in the EO771 model (P < 0.0001; **Fig. 1B, C**). Tumor growth curves showed robust suppression of tumor progression exclusively in the triple-therapy group (**Fig. 1B, C**).

**Figure 1.**
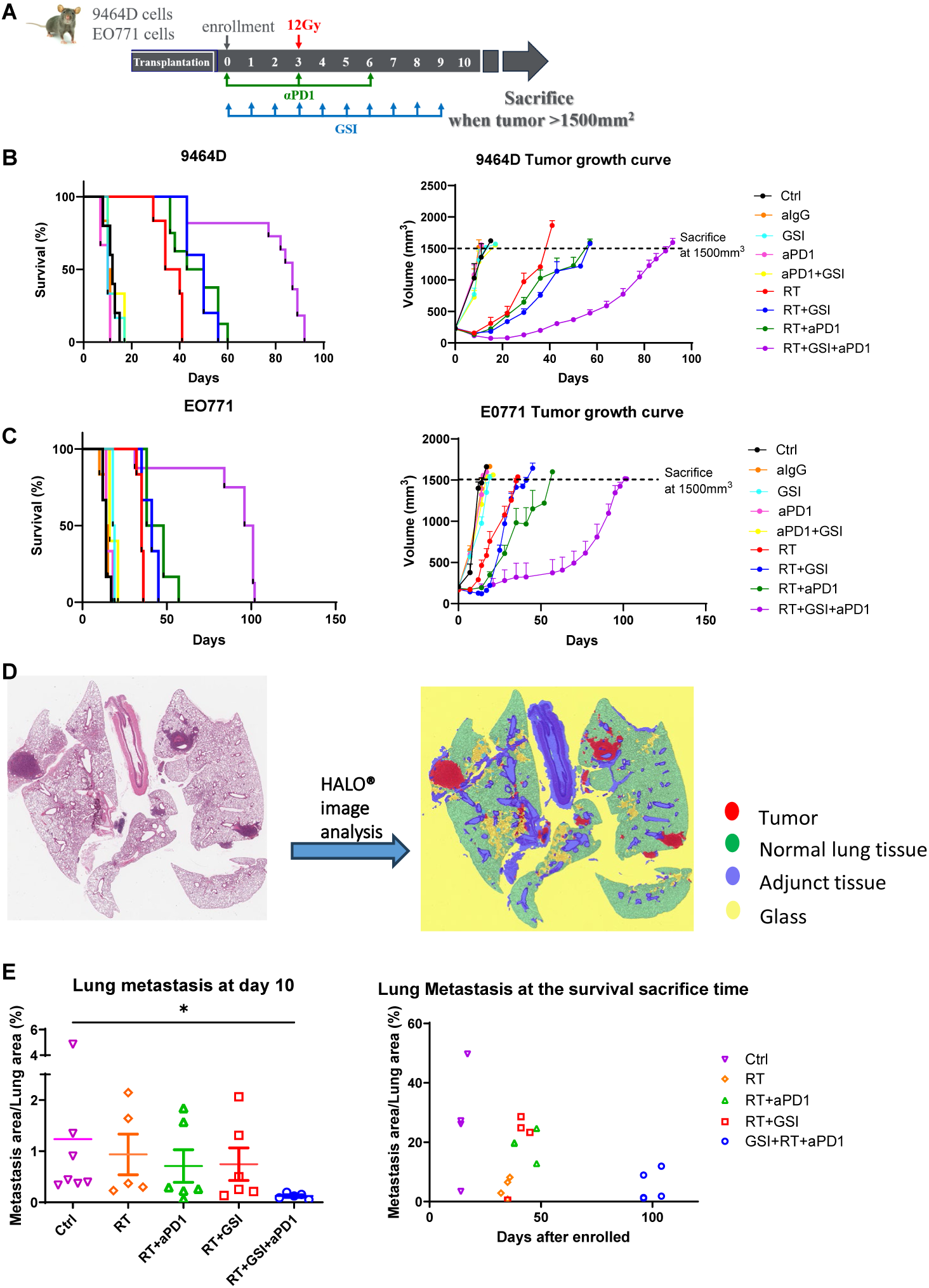
RT+aPD-1+GSI significantly prolongs survival in two separate syngeneic tumor models and reduces lung metastasis in the TNBC model. (A). Tumor cells were implanted into the flank (1 × 10^6^ 9464D) or mammary fat pad (1 × 10^5^ EO771) of C57BL/6 mice, enrolled when tumors reached 150-200mm^3^. Mice were treated as per the schema and sacrificed when tumors reached a size 1.5cm^3^. (B) The Kaplan-Meier survival curve of mice and tumor volume growth curve of 9464D tumor. Median Survival: Control 12d; IgG 10d; GSI 10d; aPD-1 10d; aPD-1+ GSI 10d; RT 37d; RT+ GSI 50d; RT+aPD-1 46d; RT+aPD-1+GSI 87d. Curve comparison by Log-rank (Mantel-Cox) test. P<0.0001. (C) The Kaplan-Meier survival curve of mice and tumor volume growth curve of EO771 tumor Median survival: Control: 14d; IgG 14.5d; GSI 18.5d; aPD-1 15d; GSI+aPD-1 10d; RT 35d; RT+ GSI 41d; RT+aPD-1 43d; RT+aPD-1+ GSI 98.5d. Curve comparison by Log-rank (Mantel-Cox) test. P<0.0001. (D) Lungs were collected from EO771 tumor–bearing mice treated as described above and sacrificed either at day 10 or at the survival endpoint. Tumor metastases were quantified using HALO image analysis, calculated as metastatic area divided by total lung area. Bars represent mean ± SD. *P<0.05, using one-way ANOVA followed by Tukey post-hoc comparisons.

To determine whether the anti-tumor effect of GSI with RT+aPD-1 treatment was mediated by a GSI-direct cytotoxic effect on tumor cells, we performed *in vitro* clonogenic survival assays testing the effect of GSI with or without radiation (**Supplementary Fig. 2**). GSI treatment did not reduce clonogenic survival or increase tumor sensitivity to RT across different radiation doses, suggesting that the treatment benefit arises through modulation of the TIME rather than inducing tumor cell death.

### Combination RT + aPD-1+GSI significantly delayed metastasis of the breast cancer tumor model

Next, we determined the effect of the triple therapy on metastasis, in the syngeneic EO771 model. GSI or aPD-1 monotherapy did not significantly reduce metastatic burden, while RT alone decreased lung metastasis compared with the non-treated control group at the survival endpoint (**Fig. 1D**). Dual-combination regimens, with either GSI or aPD-1 to RT failed to confer additional meaningful benefit. In contrast, triple therapy markedly delayed metastatic progression over RT alone. RT + aPD-1 + GSI group exhibited a significant reduction in the percentage of metastatic area relative to total lung tissue compared with non-treated control group at both the early time point (day 10; P < 0.001; **Fig. 1D**) and the survival endpoint (P < 0.001; **Fig. 1E**). Together, these findings demonstrated that triple therapy with RT + aPD-1 + GSI was required to achieve meaningful suppression of metastatic progression, with neither monotherapy nor dual-combination regimens sufficient to significantly reduced metastatic burden in the EO771 model.

### RT + aPD-1 + GSI triple therapy reprogrammed the tumor myeloid compartment toward a pro-inflammatory phenotype and induces M-MDSC expansion

To investigate how GSI influences radiation-induced innate immune responses, we first analyzed macrophage phenotypes in 9464D tumors using scRNA sequencing. The combination of RT + aPD-1 + GSI markedly altered the macrophage landscape with a shift toward a pro-inflammatory transcriptional profile (**Fig. 2A and 2B**). UMAP visualization revealed enrichment of inflammatory macrophages expressing *Il1b* and *Tnf*, whereas immunosuppressive associated genes such as *Mrc1* and *Mgl2* were reduced (**Fig. 2C**). Consistently, the M-MDSC module score (including *Arg1*, *Ly6c*, *S100a8/9*, *Il1b*, *Ptgs2*, and *Itgam*) was significantly increased following triple therapy, suggesting activation and metabolic remodeling of tumor-associated myeloid populations.

**Figure 2.**
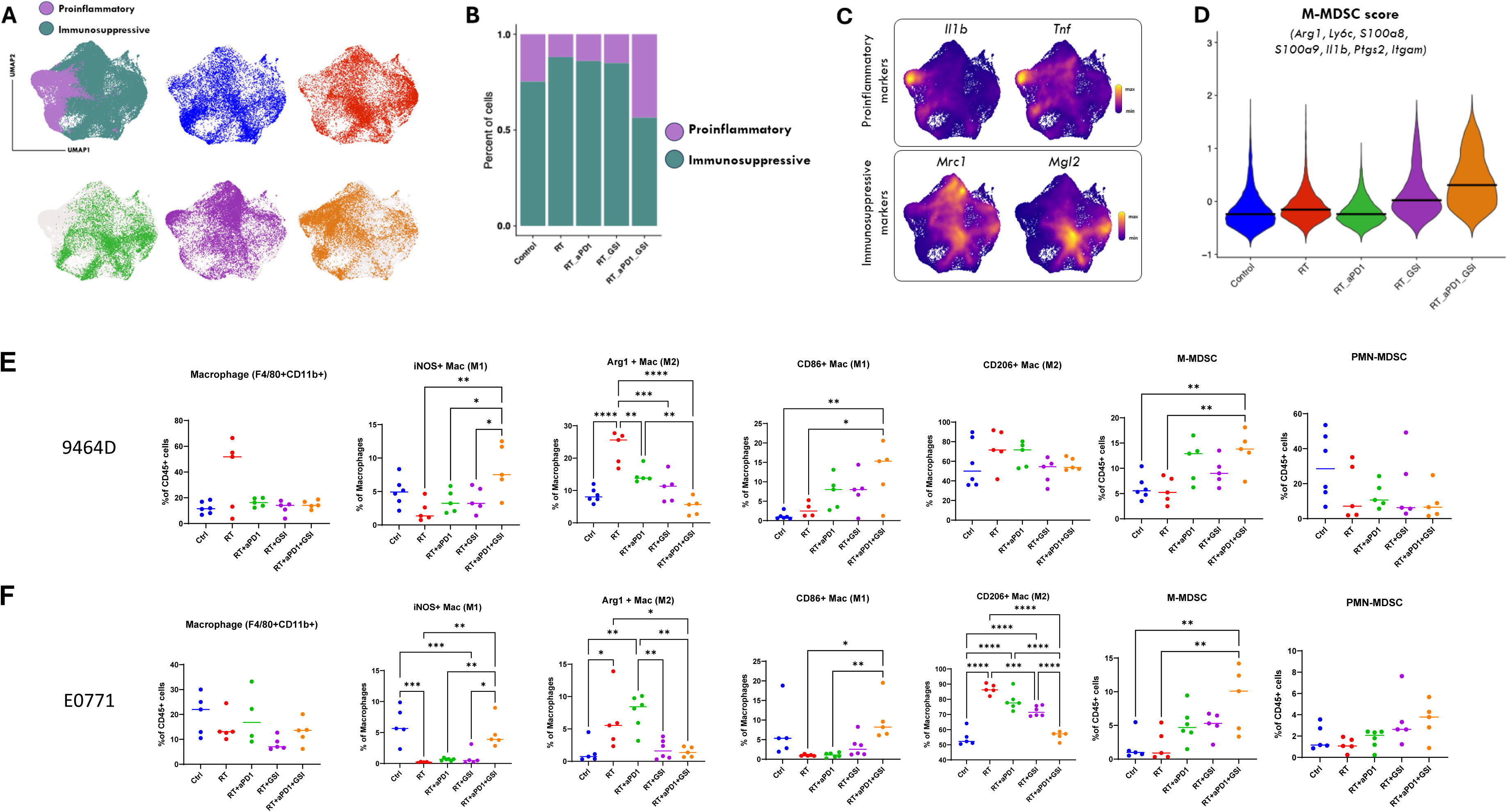
Triple therapy of RT+aPD-1+GSI enhances pro-inflammatory polarization and elevates M-MDSC scores in macrophages. (A) UMAPs showing pro-inflammatory and immunosuppressive macrophage states across treatments in 9464D tumor scRNA-seq. (B) Proportion of pro-inflammatory and immunosuppressive macrophages by treatment group (C) Expression of classical pro-inflammatory (Il1b, Tnf) and immunosuppressive (Mrc1, Mgl2) markers. (D) M-MDSC module scores increase following RT + aPD-1 + GSI (E) Changes in the proportion of pro-inflammatory and immunosuppressive macrophages subpopulation and Myeloid-derived suppressor cells (MDSC) subpopulations by flow cytometry in 9464D tumor (F) Changes in the proportion of pro-inflammatory and immunosuppressive macrophages subpopulation and MDSC subpopulations by flow cytometry in EO771 tumor. Bars represent the mean. Each dot represents one animal. *P<0.05, **P<0.01, ***P<0.001, ****P<0.0001 using one-way ANOVA followed by Tukey post-hoc comparisons.

Flow cytometry confirmed these findings in both 9464D and EO771 tumor models, showing an increase in iNOS⁺ (pro-inflammatory) macrophages and CD86⁺ macrophages, accompanied by a reduction in Arg1⁺ (immunosuppressive) macrophages (**Fig. 2E and 2F**). Notably, M-MDSC frequencies were elevated, while PMN-MDSCs remained largely unchanged (**Fig. 2E and 2F**). Together, these results show that RT +GSI +aPD-1 promote macrophage polarization to a pro-inflammatory phenotype at the expense of immunosuppressive monocyte populations, collectively reshaping the myeloid compartment in response to therapy.

### RT + aPD-1 + GSI triple therapy enhanced dendritic cell activation and antigen-presenting function despite reduced overall DC frequency

We next examined how GSI modulates dendritic cell (DC) populations and their activation state within the TME (**Fig. 3**). ScRNA sequencing of 9464D tumors revealed that RT combined with aPD-1 and GSI significantly increased expression of genes associated with DC activation and antigen presentation, including *Cd9*, *Tnfrsf9*, *Ifitm2*, *Bst2*, and proteasome components (*Psmb6*, *Psmb7*, *Psmb8*) (**Fig. 3A**). Despite a moderate reduction in the overall proportion of CD11c⁺MHC-II⁺ DCs following triple therapy, flow cytometry demonstrated a marked enrichment of the cross-presenting CD103⁺ MHC-II⁺ DC subset in both 9464D and EO771 tumor models (**Fig. 3B and 3C**). These CD103⁺ DCs were key mediators of tumor antigen presentation and CD8⁺ T cell priming, suggesting enhanced functional maturation despite reduced total DC numbers. Together, these data showed that triple therapy selectively augments the activation and differentiation of antigen-presenting DC subsets.

**Figure 3.**
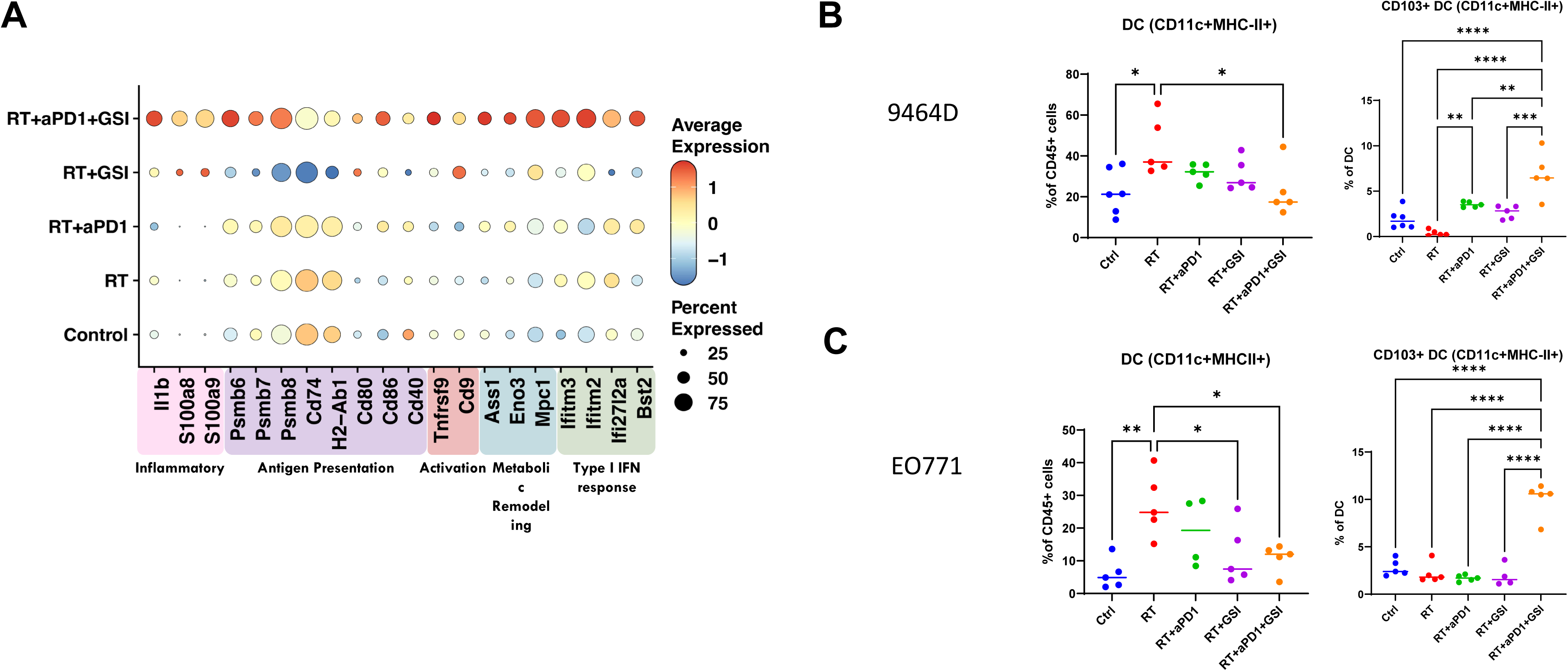
Treatment with GSI + aPD-1 reduced the overall dendritic cell (DC) proportion in RT-treated tumors, and triple-therapy of RT+aPD-1+GSI enhances DC activation and antigen-presentation. (A) Triple therapy enhances DC activation and antigen-presentation gene expression in 9464D tumor scRNA-seq. (B) Changes in the proportion of DC and CD103+DC by flowcytometry in 9464D tumor. (C) Changes in the proportion of DC and CD103+DC by flowcytometry in EO771 tumor. Bars represent the mean, each dot represents one animal. *P<0.05, **P<0.01, ***P<0.001, ****P<0.0001 using one-way ANOVA followed by Tukey post-hoc comparisons.

### RT + aPD-1 + GSI triple therapy expanded early activated CD69⁺ PD-1⁺ CD8⁺ T-cells and reduced T-cell exhaustion and Treg populations

We next examined the effects of triple therapy on the T-cell compartment (**Fig. 4**). ScRNA-seq of 9464D tumors revealed that RT + aPD-1 + GSI markedly reshaped the CD8⁺ T-cell landscape, expanding stem-like/early activated subsets (*Tcf7*⁺*Slamf6*⁺*Il7r*⁺) while contracting terminally exhausted populations (*Tox*⁺*Havcr2*⁺*Lag3*⁺). Triple therapy increased expression of genes associated with stemness and self-renewal (*Tcf7*, *Ccr7*, *Il7r*, *Lef1*), along with upregulation of type I interferon response genes (*Ifitm2*, *Bst2*, *Mx1*, *Isg15*) (**Fig. 4A-E**). Consistent with the scRNA-seq data, flow cytometry analysis demonstrated that triple therapy significantly increased the number of early activated CD69⁺ PD-1⁺ CD8⁺ T cells. In addition, triple therapy significantly reduced both terminally exhausted TCF1⁻Tim3⁺ CD8⁺ T cells and progenitor exhausted TCF1⁺Tim3⁻ CD8⁺ T subsets compared with RT + aPD-1 in 9464D tumors (**Fig 4. F-G**). Moreover, regulatory T cells (CD4⁺FoxP3⁺CD25⁺) were substantially decreased, whereas NK1.1⁺ NK cells were increased in the triple therapy group, indicating a shift from an immunosuppressive T-cell-dominated microenvironment toward broader activation of cytotoxic T-cell populations.

**Figure 4.**
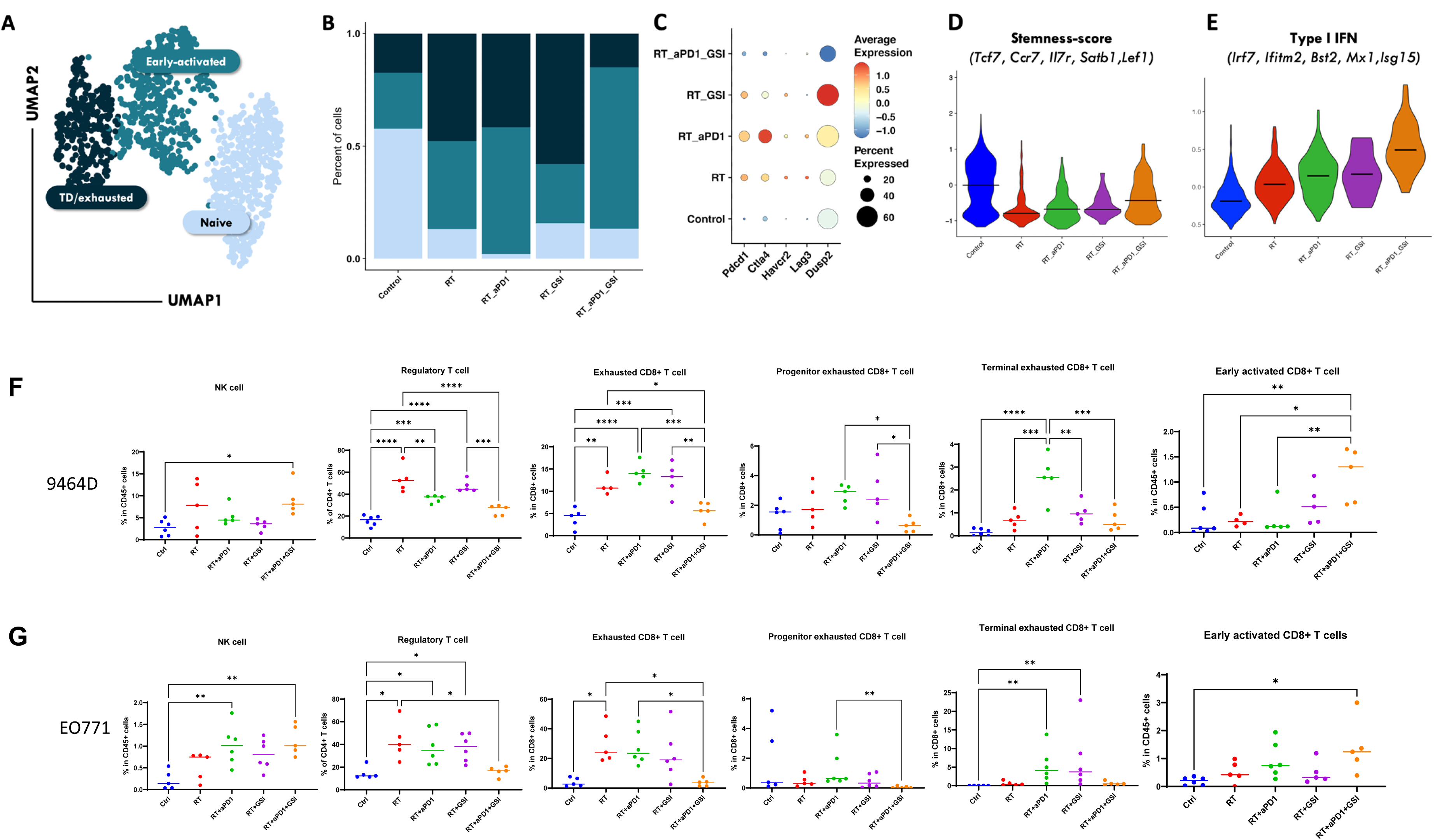
Triple therapy of RT+GSI+aPD-1 expands early-activated CD8⁺ T cells while reducing exhaustion and regulatory T cells. (A–B). UMAP and subset composition showing increased early-activated and decreased terminally exhausted CD8⁺ T cells after triple therapy in 9464D tumor scRNAseq. (C). Expression of canonical exhaustion and checkpoint genes across treatment groups (D–E). Triple therapy enhances CD8⁺ stemness and type I IFN module scores (F-G). Proportion changes of NK1.1+ NK cell, CD4+FoxP3+CD25+ Treg cells, PD-1+CD69- exhausted CD8+ T cells, progenitor exhausted TCF1^+^Tim3^-^ CD8 T cells, terminal exhausted TCF1⁻Tim3⁺ CD8 T cells and early active CD69+ PD-1+ CD8+ T cells by flowcytometry in 9464D tumor and E0771 tumor. Bars represent mean, each dot represents one animal. *P<0.05, **P<0.01, ***P<0.001, ****P<0.0001 using one-way ANOVA followed by Tukey post-hoc comparisons.

### Lack of anti-tumor effect of RT + aPD-1 + GSI in immunodeficient nude and macrophage-depleted mice

To determine whether the therapeutic efficacy of the triple combination depends on an intact adaptive immune system, we implanted 9464D (subcutaneously) and EO771 cells (fat pad) into athymic nude mice, which lack functional T cells, and treated them according to the schema in **Fig. 1A**. In contrast to the results in immunocompetent mice, the combination of RT+ aPD-1+ GSI did not significantly improve tumor control or mice survival in either model. Tumor growth curves showed comparable kinetics among treatment groups, and Kaplan–Meier analyses revealed no significant survival benefit following triple therapy (**Fig. 5A and B vs Fig. 1 B and C**). Similarly, in the EO771 model, lung metastasis burden remained unchanged among treated and control groups at the survival endpoint (**Fig 5B**).

**Figure 5.**
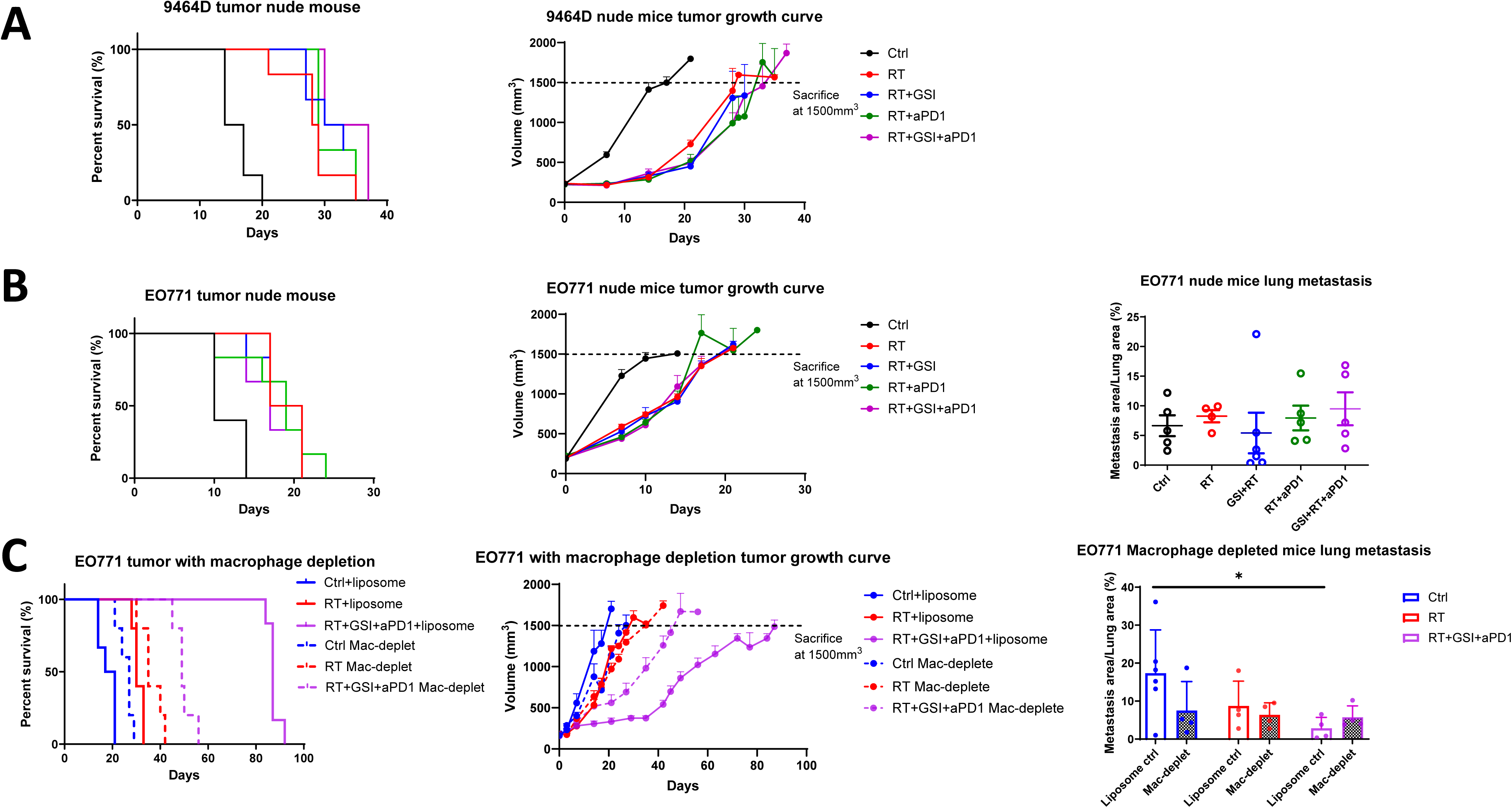
Combination treatment of RT+GSI+aPD-1 does not inhibit tumors in athymic nude mice and macrophage-depleted mice. 9464D cells or EO771 were implanted into nude mice and macrophage-depleted followed by treatment per the schema in Fig. 1. Mice were sacrificed when tumors reached a size >1.5 cm^3^. (A). The Kaplan-Meier survival curve of mice and tumor volume growth curve of athymic nude mice 9464D tumor. Median Survival: Control: 15.5d; RT 28.5d; RT+ GSI 28.5d; RT+aPD-1 29d; RT+aPD-1+ GSI 30d. Curve comparison by Log-rank (Mantel-Cox) test. p>0.05. (B). The Kaplan-Meier survival curve of mice and tumor volume growth curve, as well as lung metastasis of athymic nude mice EO771 tumor at the survival endpoint. The Kaplan-Meier survival curve of mice and tumor volume growth curve for EO771. Median Survival: Control: 10d; RT 19d; RT+ GSI 19d; RT+aPD-1 19d; RT+aPD-1+ GSI 17d. Curve comparison by Log-rank (Mantel-Cox) test. P>0.05. (C). The Kaplan-Meier survival curve of mice and tumor volume growth curve, as well as lung metastasis of macrophage depleted in EO771 tumor mice at the survival endpoint. Liposome control: Control: 17d; RT 28d; RT+aPD-1+ GSI 87d. Macrophage depleted: Control: 27d; RT 35d; RT+aPD-1+ GSI 49d. Comparison by Log-rank (Mantel-Cox) test and one-way ANOVA.

To further evaluate the contribution of macrophages, we performed macrophage depletion using clodronate liposomes in the syngeneic EO771 model in control, RT and triple therapy group **(Fig. 5C)**. Consistent with literatures^26, 27^, macrophage depletion prolonged survival in both the untreated control (p<0.01) and RT-alone groups (p<0.05). In contrast, macrophage-depleted mice exhibited shorter survival with triple therapy when compared with liposome controlled triple therapy group (p<0.01). In liposome control mice, triple therapy reduced lung metastasis compared with the untreated control group at the survival endpoint (p<0.01), whereas in clodronate-treated macrophage-depleted mice, metastatic burden was comparable across all treatment groups (**Fig. 5C**). There was an increase in the triple therapy group in lung metastasis with clodronate treatment compared to liposomal control, however, it was not statistically significant.

Together, these findings indicated that the enhanced anti-tumor efficacy of RT + aPD-1 + GSI observed in immunocompetent mice is dependent on immune cell activity, particularly T-cell and macrophage-mediated mechanisms.

## Discussion

By targeting gamma-secretase with the GSI AL101 in combination with RT and aPD-1 therapy, we observed significant suppression of primary tumor growth and metastasis and more than doubled time of survival compared with the radiation-treated groups. Importantly, therapeutic efficacy was observed in both an immunologically warm, partially ICI-sensitive model (EO771 triple-negative breast cancer) and a cold, ICI-resistant model (9464D neuroblastoma; low TMB, minimal T cell infiltration), underscoring the broad applicability of RT + aPD-1 + GSI triple therapy across distinct immune contextures.. Recent clinical progress also supported the translational feasibility of GSIs: nirogacestat received FDA approval for desmoid tumors following the phase 3 DeFi trial^28^, while AL101/AL102^29, 30^ and CB-103^31^ showed early promise adenoid cystic carcinoma and desmoid tumors with Notch-activation. These advances highlight the potential to combine GSIs with RT and ICI across tumor types. GSI is a distinct proteolytic complex required for the activation of many transmembrane proteins. The cleavage of substrates by γ-secretase plays diverse biological roles in producing essential products for the organism. More than 90 transmembrane proteins have been reported to be substrates of γ-secretase. The most well-known is Notch pathway^32, 33^.

Here we found that GSI drives robust immune reprogramming in two-immunogenic phenotypes tumor models, with macrophage depletion fully abolishing the ability of the GSI to improve the aPD-1 and RT regimen. Triple therapy repolarized TAMs toward a pro-inflammatory phenotype while reducing immunosuppressive myeloid populations. We propose that this occurs through GSI inhibition of Notch signaling which is consistent with prior reports that Notch inhibition remodels the myeloid TME^20, 34^. For example, genetic Notch inhibition in myeloid cells reduced tumor growth and macrophage infiltration in pancreatic cancer^35^ ^36^. Interestingly, recent studies have shown that GSI in combination with aPD-1 modulate antitumor immunity by decreasing macrophage infiltration and reducing PD-L1 expression at lung metastatic sites in TNBC models, a process mediated in part by CCL2 and IL-1β^20^.

Despite broad myeloid reprogramming, M-MDSCs emerged as a compensatory suppressive population. Previous studies have also shown that MDSCs can promote tumor progression by enhancing cancer stem-like properties and suppressing T-cell activation, in part through Notch-dependent signaling pathways^37, 38^. Therapy-induced inflammatory cues, including signaling through CSF1R^39^ or CXCR2^40^, may drive M-MDSC activation or accumulation in this context. Together, these findings suggest that M-MDSCs may represent a potential target for additional combinatorial therapeutic strategies^41^ ^42^.

Although total DC numbers declined with triple therapy, the remaining DCs - particularly CD103⁺ DCs, show enhanced activation and cross-presentation, suggesting that GSI amplifies RT-induced immunogenicity by promoting DC maturation and antigen presentation. This finding aligns with evidence that Notch restrains DC development and type I IFN activation, and that Notch inhibition enhances T-cell cross-priming^43^. Notably, Notch-mediated DC-T cell communication can be bidirectional, with Jagged-1 and Jagged-2 promoting Treg differentiation and function^44, 45^.

We observed that the GSI on top of the RT and aPD1 could modulate the T-cell activation and exhaustion. Controversy remains regarding the contribution of Notch signaling to T-cell activity and tumorigenesis and this maybe context dependent or tumor type dependent^46^. Notch1 loss in recurrent laryngeal cancer tissue sample was associated with reduced TIL infiltration and immunosurveillance escape^47^. Notch inhibition reduced PD-1 expression on CD8⁺ T cells in colorectal carcinoma, suggesting that Notch can suppress T-cell function^48^. Additionally, T-cell senescence has been correlated with Notch1 intracellular domain expression^49^. We found that triple therapy expanded early activated CD69⁺ PD-1⁺ CD8⁺ T cells and progenitor-like CD8⁺ T cell populations while reducing terminal exhaustion, suggesting GSI enhances T cell priming and preserves effector potential by limiting exhaustion and Treg-mediated suppression. Critically, this efficacy was fully abrogated in athymic nude mice, confirming that T cell-mediated adaptive immunity is indispensable for the tumor control and metastasis suppression observed with this combination.

Several limitations in our study should be acknowledged. First, because GSIs affect not only Notch signaling but multiple pathways, the safety, timing, and dosing, as well as optimal radiation regimens, require careful evaluation in clinically relevant models. Second, GSIs act as pan-Notch suppressor by blocking receptor cleavage, and further mechanistic studies are needed to define the specific Notch pathways involved.

Our findings support GSI as a strategy to enhance systemic antitumor immunity and abscopal responses. With Notch inhibitors like AL101 already in trials^50^, this work establishes a mechanistically grounded strategy to overcome radio-immunotherapy resistance and will generate data to support the development of future clinical trials.

## Acknowledgements

This research was funded in part by the NIH/NCI Cancer Center Support Grant P30CA013696 (Oncology Precision Therapeutics and Imaging Core Shared Resource). This work was supported by the Velocity Grant and Research Stabilization Funding from Columbia University Irving Medical Center. We thank the Institute of Comparative Medicine (ICM) Animal Facility, the Columbia Stem Cell Initiative (CSCI) Flow Cytometry Core, the Systems Biology Single-Cell Core, and the Digital and Computational Pathology Lab (DCPL) shared resource at Columbia University Irving Medical Center for their technical support and expertise.

## References

1. Machiraju D, Schäfer S, Hassel JC. Potential Reasons for Unresponsiveness to Anti-PD1 Immunotherapy in Young Patients with Advanced Melanoma. Life (Basel). 2021;11(12). Epub 20211130. doi: 10.3390/life11121318. PubMed PMID: 34947849; PMCID: PMC8707626.

2. Huang AC, Postow MA, Orlowski RJ, Mick R, Bengsch B, Manne S, Xu W, Harmon S, Giles JR, Wenz B. T-cell invigoration to tumour burden ratio associated with anti-PD-1 response. Nature. 2017;545(7652):60–5.

3. Zhang Z, Liu X, Chen D, Yu J. Radiotherapy combined with immunotherapy: the dawn of cancer treatment. Signal Transduction and Targeted Therapy. 2022;7(1):258. doi: 10.1038/s41392-022-01102-y.

4. Liao Y, Deng J, Yang X, Wang D, Du X. Advances in radiotherapy enhancing the efficacy of immune checkpoint inhibitors in malignant. Front Oncol. 2025;15:1611036. Epub 20250701. doi: 10.3389/fonc.2025.1611036. PubMed PMID: 40666094; PMCID: PMC12259436.

5. Muroyama Y, Nirschl TR, Kochel CM, Lopez-Bujanda Z, Theodros D, Mao W, Carrera-Haro MA, Ghasemzadeh A, Marciscano AE, Velarde E, Tam AJ, Thoburn CJ, Uddin M, Meeker AK, Anders RA, Pardoll DM, Drake CG. Stereotactic Radiotherapy Increases Functionally Suppressive Regulatory T Cells in the Tumor Microenvironment. Cancer Immunol Res. 2017;5(11):992–1004. Epub 20171002. doi: 10.1158/2326-6066.CIR-17-0040. PubMed PMID: 28970196; PMCID: PMC5793220.

6. Shi X, Shiao SL. The role of macrophage phenotype in regulating the response to radiation therapy. Translational Research. 2018;191:64–80.

7. Ma R-Y, Black A, Qian B-Z. Macrophage diversity in cancer revisited in the era of single-cell omics. Trends in Immunology. 2022.

8. Hartley F, Ebert M, Cook AM. Leveraging radiotherapy to improve immunotherapy outcomes: rationale, progress and research priorities. Clin Transl Immunology. 2025;14(4):e70030. Epub 20250408. doi: 10.1002/cti2.70030. PubMed PMID: 40206193; PMCID: PMC11977402.

9. Andersson ER, Sandberg R, Lendahl U. Notch signaling: simplicity in design, versatility in function. Development. 2011;138(17):3593–612.

10. Katoh M, Katoh M. Integrative genomic analyses on HES/HEY family: Notch-independent HES1, HES3 transcription in undifferentiated ES cells, and Notch-dependent HES1, HES5, HEY1, HEY2, HEYL transcription in fetal tissues, adult tissues, or cancer. International journal of oncology. 2007;31(2):461–6.

11. Shi Q, Xue C, Zeng Y, Yuan X, Chu Q, Jiang S, Wang J, Zhang Y, Zhu D, Li L. Notch signaling pathway in cancer: from mechanistic insights to targeted therapies. Signal Transduction and Targeted Therapy. 2024;9(1):128. doi: 10.1038/s41392-024-01828-x.

12. Amsen D, Helbig C, Backer RA. Notch in T cell differentiation: all things considered. Trends in immunology. 2015;36(12):802–14.

13. Samon JB, Champhekar A, Minter LM, Telfer JC, Miele L, Fauq A, Das P, Golde TE, Osborne BA. Notch1 and TGFβ1 cooperatively regulate Foxp3 expression and the maintenance of peripheral regulatory T cells. Blood, The Journal of the American Society of Hematology. 2008;112(5):1813–21.

14. Li X, Yan X, Wang Y, Kaur B, Han H, Yu J. The Notch signaling pathway: a potential target for cancer immunotherapy. J Hematol Oncol. 2023;16(1):45. Epub 2023/05/03. doi: 10.1186/s13045-023-01439-z. PubMed PMID: 37131214; PMCID: PMC10155406.

15. Lin Y, Zhao J-L, Zheng Q-J, Jiang X, Tian J, Liang S-Q, Guo H-W, Qin H-Y, Liang Y-M, Han H. Notch signaling modulates macrophage polarization and phagocytosis through direct suppression of signal regulatory protein α expression. Frontiers in immunology. 2018;9:1744.

16. Meurette O, Mehlen P. Notch signaling in the tumor microenvironment. Cancer cell. 2018;34(4):536–48.

17. Cahill EF, Tobin LM, Carty F, Mahon BP, English K. Jagged-1 is required for the expansion of CD4+ CD25+ FoxP3+ regulatory T cells and tolerogenic dendritic cells by murine mesenchymal stromal cells. Stem cell research & therapy. 2015;6(1):1–13.

18. Liu H, Wang J, Zhang M, Xuan Q, Wang Z, Lian X, Zhang Q. Jagged1 promotes aromatase inhibitor resistance by modulating tumor-associated macrophage differentiation in breast cancer patients. Breast Cancer Res Treat. 2017;166(1):95–107. Epub 20170720. doi: 10.1007/s10549-017-4394-2. PubMed PMID: 28730338.

19. Wang YC, He F, Feng F, Liu XW, Dong GY, Qin HY, Hu XB, Zheng MH, Liang L, Feng L, Liang YM, Han H. Notch signaling determines the M1 versus M2 polarization of macrophages in antitumor immune responses. Cancer Res. 2010;70(12):4840–9. Epub 20100525. doi: 10.1158/0008-5472.Can-10-0269. PubMed PMID: 20501839.

20. Shen Q, Murakami K, Sotov V, Butler M, Ohashi PS, Reedijk M. Inhibition of Notch enhances efficacy of immune checkpoint blockade in triple-negative breast cancer. Science Advances. 2024;10(44):eado8275. doi: doi:10.1126/sciadv.ado8275.

21. McCaw TR, Inga E, Chen H, Jaskula-Sztul R, Dudeja V, Bibb JA, Ren B, Rose JB. Gamma Secretase Inhibitors in Cancer: A Current Perspective on Clinical Performance. Oncologist. 2021;26(4):e608–e21. Epub 20210102. doi: 10.1002/onco.13627. PubMed PMID: 33284507; PMCID: PMC8018325.

22. Banerjee D, Barton SM, Grabham PW, Rumeld AL, Okochi S, Street C, Kadenhe-Chiweshe A, Boboila S, Yamashiro DJ, Connolly EP. High-Dose Radiation Increases Notch1 in Tumor Vasculature. International Journal of Radiation Oncology*Biology*Physics. 2020;106(4):857–66. doi: 10.1016/j.ijrobp.2019.11.010.

23. Franken NA, Rodermond HM, Stap J, Haveman J, van Bree C. Clonogenic assay of cells in vitro. Nat Protoc. 2006;1(5):2315–9. doi: 10.1038/nprot.2006.339. PubMed PMID: 17406473.

24. Boboila S, Okochi S, Banerjee D, Barton S, Street C, Zenilman AL, Wang Q, Gartrell RD, Saenger YM, Welch D, Wu CC, Kadenhe-Chiweshe A, Yamashiro DJ, Connolly EP. Combining immunotherapy with high-dose radiation therapy (HDRT) significantly inhibits tumor growth in a syngeneic mouse model of high-risk neuroblastoma. Heliyon. 2023;9(6):e17399. Epub 20230619. doi: 10.1016/j.heliyon.2023.e17399. PubMed PMID: 37408891; PMCID: PMC10319189.

25. Boboila S, Okochi S, Banerjee D, Barton S, Street C, Zenilman AL, Wang Q, Gartrell RD, Saenger YM, Welch D, Wu C-C, Kadenhe-Chiweshe A, Yamashiro DJ, Connolly EP. Combining immunotherapy with high-dose radiation therapy (HDRT) significantly inhibits tumor growth in a syngeneic mouse model of high-risk neuroblastoma. Heliyon. 2023;9(6):e17399. doi: 10.1016/j.heliyon.2023.e17399.

26. Hamon P, Gerbé De Thoré M, Classe M, Signolle N, Liu W, Bawa O, Meziani L, Clémenson C, Milliat F, Deutsch E, Mondini M. TGFβ receptor inhibition unleashes interferon-β production by tumor-associated macrophages and enhances radiotherapy efficacy. J Immunother Cancer. 2022;10(3). doi: 10.1136/jitc-2021-003519. PubMed PMID: 35301235; PMCID: PMC8932273.

27. Genard G, Lucas S, Michiels C. Reprogramming of Tumor-Associated Macrophages with Anticancer Therapies: Radiotherapy versus Chemo- and Immunotherapies. Front Immunol. 2017;8:828. Epub 20170714. doi: 10.3389/fimmu.2017.00828. PubMed PMID: 28769933; PMCID: PMC5509958.

28. Gounder M, Ratan R, Alcindor T, Schöffski P, van der Graaf WT, Wilky BA, Riedel RF, Lim A, Smith LM, Moody S, Attia S, Chawla S, D’Amato G, Federman N, Merriam P, Van Tine BA, Vincenzi B, Benson C, Bui NQ, Chugh R, Tinoco G, Charlson J, Dileo P, Hartner L, Lapeire L, Mazzeo F, Palmerini E, Reichardt P, Stacchiotti S, Bailey HH, Burgess MA, Cote GM, Davis LE, Deshpande H, Gelderblom H, Grignani G, Loggers E, Philip T, Pressey JG, Kummar S, Kasper B. Nirogacestat, a γ-Secretase Inhibitor for Desmoid Tumors. N Engl J Med. 2023;388(10):898–912. doi: 10.1056/NEJMoa2210140. PubMed PMID: 36884323; PMCID: PMC11225596.

29. Ferrarotto R, Mishra V, Herz E, Yaacov A, Solomon O, Rauch R, Mondshine A, Motin M, Leibovich-Rivkin T, Davis M, Kaye J, Weber CR, Shen L, Pearson AT, Rosenberg AJ, Chen X, Singh A, Aster JC, Agrawal N, Izumchenko E. AL101, a gamma-secretase inhibitor, has potent antitumor activity against adenoid cystic carcinoma with activated NOTCH signaling. Cell Death Dis. 2022;13(8):678. Epub 20220805. doi: 10.1038/s41419-022-05133-9. PubMed PMID: 35931701; PMCID: PMC9355983.

30. Gounder MM, Jones RL, Chugh R, Agulnik M, Singh AS, Tine BAV, Andelkovic V, Choy E, Lewin JH, Ratan R, Gordon GB, Yovell J, Gutierrez AA, Kasper B. RINGSIDE phase 2/3 trial of AL102 for treatment of desmoid tumors (DT): Phase 2 results. Journal of Clinical Oncology. 2023;41(16_suppl):11515–. doi: 10.1200/JCO.2023.41.16_suppl.11515.

31. Hanna GJ, Stathis A, Lopez-Miranda E, Racca F, Quon D, Leyvraz S, Hess D, Keam B, Rodon J, Ahn MJ, Kim HR, Schneeweiss A, Ribera JM, DeAngelo D, Perez Garcia JM, Cortes J, Schönborn-Kellenberger O, Weber D, Pisa P, Bauer M, Beni L, Bobadilla M, Lehal R, Vigolo M, Vogl FD, Garralda E. A Phase I Study of the Pan-Notch Inhibitor CB-103 for Patients with Advanced Adenoid Cystic Carcinoma and Other Tumors. Cancer Res Commun. 2023;3(9):1853–61. doi: 10.1158/2767-9764.Crc-23-0333. PubMed PMID: 37712875; PMCID: PMC10501326.

32. Cheng Y-L, Choi Y, Sobey CG, Arumugam TV, Jo D-G. Emerging roles of the γ-secretase-notch axis in inflammation. Pharmacology & Therapeutics. 2015;147:80–90. doi: 10.1016/j.pharmthera.2014.11.005.

33. Doerfler P, Shearman MS, Perlmutter RM. Presenilin-dependent γ-secretase activity modulates thymocyte development. Proceedings of the National Academy of Sciences. 2001;98(16):9312–7. doi: doi:10.1073/pnas.161102498.

34. Wang M, Yu F, Zhang Y, Li P. Novel insights into Notch signaling in tumor immunity: potential targets for cancer immunotherapy. Frontiers in Immunology. 2024;15:1352484.

35. Yan W, Menjivar RE, Bonilla ME, Steele NG, Kemp SB, Du W, Donahue KL, Brown KL, Carpenter ES, Avritt FR, Irizarry-Negron VM, Yang S, Burns WR, 3rd, Zhang Y, Pasca di Magliano M, Bednar F. Notch Signaling Regulates Immunosuppressive Tumor-Associated Macrophage Function in Pancreatic Cancer. Cancer Immunol Res. 2024;12(1):91–106. doi: 10.1158/2326-6066.Cir-23-0037. PubMed PMID: 37931247; PMCID: PMC10842043.

36. Geng Y, Fan J, Chen L, Zhang C, Qu C, Qian L, Chen K, Meng Z, Chen Z, Wang P. A Notch-Dependent Inflammatory Feedback Circuit between Macrophages and Cancer Cells Regulates Pancreatic Cancer Metastasis. Cancer Res. 2021;81(1):64–76. Epub 20201110. doi: 10.1158/0008-5472.Can-20-0256. PubMed PMID: 33172931.

37. Peng D, Tanikawa T, Li W, Zhao L, Vatan L, Szeliga W, Wan S, Wei S, Wang Y, Liu Y, Staroslawska E, Szubstarski F, Rolinski J, Grywalska E, Stanisławek A, Polkowski W, Kurylcio A, Kleer C, Chang AE, Wicha M, Sabel M, Zou W, Kryczek I. Myeloid-Derived Suppressor Cells Endow Stem-like Qualities to Breast Cancer Cells through IL6/STAT3 and NO/NOTCH Cross-talk Signaling. Cancer Res. 2016;76(11):3156–65. Epub 20160406. doi: 10.1158/0008-5472.Can-15-2528. PubMed PMID: 27197152; PMCID: PMC4891237.

38. Shi H, Zhu Y, Shang K, Tian T, Yin Z, Shi J, He Y, Ding J, Zhang F. The role of notch signaling in regulating myeloid-derived suppressor cells: Implications in Cancer and autoimmune diseases. Int Immunopharmacol. 2025;157:114693. Epub 20250429. doi: 10.1016/j.intimp.2025.114693. PubMed PMID: 40306114.

39. Xu J, Escamilla J, Mok S, David J, Priceman S, West B, Bollag G, McBride W, Wu L. CSF1R signaling blockade stanches tumor-infiltrating myeloid cells and improves the efficacy of radiotherapy in prostate cancer. Cancer Res. 2013;73(9):2782–94. Epub 20130215. doi: 10.1158/0008-5472.Can-12-3981. PubMed PMID: 23418320; PMCID: PMC4097014.

40. Kumar V, Patel S, Tcyganov E, Gabrilovich DI. The Nature of Myeloid-Derived Suppressor Cells in the Tumor Microenvironment. Trends Immunol. 2016;37(3):208–20. Epub 20160206. doi: 10.1016/j.it.2016.01.004. PubMed PMID: 26858199; PMCID: PMC4775398.

41. Veglia F, Sanseviero E, Gabrilovich DI. Myeloid-derived suppressor cells in the era of increasing myeloid cell diversity. Nature Reviews Immunology. 2021;21(8):485–98. doi: 10.1038/s41577-020-00490-y.

42. Veglia F, Sanseviero E, Gabrilovich DI. Myeloid-derived suppressor cells in the era of increasing myeloid cell diversity. Nat Rev Immunol. 2021;21(8):485–98. Epub 20210201. doi: 10.1038/s41577-020-00490-y. PubMed PMID: 33526920; PMCID: PMC7849958.

43. Campbell DJ, Ziegler SF. FOXP3 modifies the phenotypic and functional properties of regulatory T cells. Nat Rev Immunol. 2007;7(4):305–10. doi: 10.1038/nri2061. PubMed PMID: 17380159.

44. Cahill EF, Tobin LM, Carty F, Mahon BP, English K. Jagged-1 is required for the expansion of CD4+ CD25+ FoxP3+ regulatory T cells and tolerogenic dendritic cells by murine mesenchymal stromal cells. Stem Cell Res Ther. 2015;6(1):19. Epub 20150311. doi: 10.1186/s13287-015-0021-5. PubMed PMID: 25890330; PMCID: PMC4414370.

45. Kared H, Adle-Biassette H, Foïs E, Masson A, Bach JF, Chatenoud L, Schneider E, Zavala F. Jagged2-expressing hematopoietic progenitors promote regulatory T cell expansion in the periphery through notch signaling. Immunity. 2006;25(5):823–34. Epub 20061102. doi: 10.1016/j.immuni.2006.09.008. PubMed PMID: 17081781.

46. Sultana J, Choudhury PR, Bera S, Chakravarti M, Guha A, Das P, Das J, Iyer GS, Sarkar A, Dhar S, Ganguly N, Baral R, Bose A, Banerjee S. Notch signalling in T cells: bridging tumour immunity and intratumoral cellular crosstalk. Front Immunol. 2025;16:1659614. Epub 20251002. doi: 10.3389/fimmu.2025.1659614. PubMed PMID: 41112285; PMCID: PMC12528159.

47. Gong XY, Chen HB, Zhang LQ, Chen DS, Li W, Chen DH, Xu J, Zhou H, Zhao LL, Song YJ, Xiao MZ, Deng WL, Qi C, Wang XR, Chen X. NOTCH1 mutation associates with impaired immune response and decreased relapse-free survival in patients with resected T1-2N0 laryngeal cancer. Front Immunol. 2022;13:920253. Epub 20220715. doi: 10.3389/fimmu.2022.920253. PubMed PMID: 35911687; PMCID: PMC9336464.

48. Yu W, Wang Y, Guo P. Notch signaling pathway dampens tumor-infiltrating CD8(+) T cells activity in patients with colorectal carcinoma. Biomed Pharmacother. 2018;97:535–42. Epub 20171106. doi: 10.1016/j.biopha.2017.10.143. PubMed PMID: 29096354.

49. Hoare M, Ito Y, Kang TW, Weekes MP, Matheson NJ, Patten DA, Shetty S, Parry AJ, Menon S, Salama R, Antrobus R, Tomimatsu K, Howat W, Lehner PJ, Zender L, Narita M. NOTCH1 mediates a switch between two distinct secretomes during senescence. Nat Cell Biol. 2016;18(9):979–92. Epub 20160815. doi: 10.1038/ncb3397. PubMed PMID: 27525720; PMCID: PMC5008465.

50. Ferrarotto R, Mishra V, Herz E, Yaacov A, Solomon O, Rauch R, Mondshine A, Motin M, Leibovich-Rivkin T, Davis M, Kaye J, Weber CR, Shen L, Pearson AT, Rosenberg AJ, Chen X, Singh A, Aster JC, Agrawal N, Izumchenko E. AL101, a gamma-secretase inhibitor, has potent antitumor activity against adenoid cystic carcinoma with activated NOTCH signaling. Cell Death & Disease. 2022;13(8):678. doi: 10.1038/s41419-022-05133-9.

