## Supplementary table1-2 and supplementary figure 1-2 for "A Novel Therapeutic Approach: Gamma Secretase Inhibitor Enhances Radiotherapy and Checkpoint Blockade Therapy via Reprogramming of the Tumor Microenvironment"

**Supplementary Table 1. T cell panel for flow cytometry**

| **Marker** | **Clone** | **Vendor** | **Cat#** |
| --- | --- | --- | --- |
| Zombie NIR |  | Biolegend | 423106 |
| CD45 BUV805 | 30-F11 | BD Horizon™ | 568336 |
| CD3e-V450 | 500A2 | BD-bioscience | 560801 |
| CD8 R718 | 53-6.7 | BD Horizon™ | 566985 |
| CD4-FITC | RM4-5 | Thermo Fisher-eBioscience | 11-0042-82 |
| CD25-BV650 | PC61 | Biolegend | 102038 |
| PD-1 PE-Cy5 | 29F.1A12 | Biolegend | 135256 |
| NK1.1-BV785 | PK136 | Biolegend | 108749 |
| TIM3 RB780 | 25F.1D6 | BD Horizon™ | 569810 |
| CD69 RB613 | H1.2F3 | BD OptiBuild™ | 758234 |
| CTLA4 BV605 | 369610 | Biolegend | 369606 |
| FOXP3-PE (intracellular) | FJK-16s | Thermo Fisher-eBioscience | 12-5773-82 |
| IFN-g APC-Fire 750 (intracellular) | XMG1.2 | Biolegend | 505859 |
| TCF-1 AF647 (intracellular) | S33-966 | BD Pharmingen™ | 566693 |
| Ki-67 RB705 (intracellular) | B56 | BD Horizon™ | 570280 |

**Supplementary Table 2. Myeloid cell panel antibodies for flow cytometry**

| **Marker** | **Clone** | **Vendor** | **Cat#** |
| --- | --- | --- | --- |
| Zombie NIR |  | Biolegend | 423106 |
| CD45 BUV805 | 30-F11 | BD Horizon™ | 568336 |
| F4/80-PerCp-Cy5.5 | BM8 | Biolegend | 123128 |
| CD8 R718 | 53-6.7 | BD Horizon™ | 566985 |
| Ly6G-APC-Fire 750 | 1A8 | Biolegend | 127652 |
| CD11b-PE | M1/70 | BD | 557397 |
| Ly6C-RB545 | AL-21 | Biolegend | 128015 |
| PD-L1-RB744 | MIH5 | BD | 758169 |
| CD206-BV650 | C068C2 | Biolegend | 385449 |
| CD86-RB780 | PO3 | BD | 755499 |
| CD11c-BV421 | N418 | Biolegend | 113312747 |
| MHC-II-FITC | M5/114.15.2 | Biolegend | 107605 |
| CD70-BUV661 | FR70 | BD | 741564 |
| XCR1-SR718 | ZET | Biolegend | 285190 |
| CD103 RB613 | M290 | BD | 758900 |
| INOS-FITC (intracellular) | CXNFT | Thermo Fisher-eBioscience | 53-5920-82 |
| Arg-1-eflour450 (intracellular) | A1exF5 | Invivogen | 48-3697-82 |

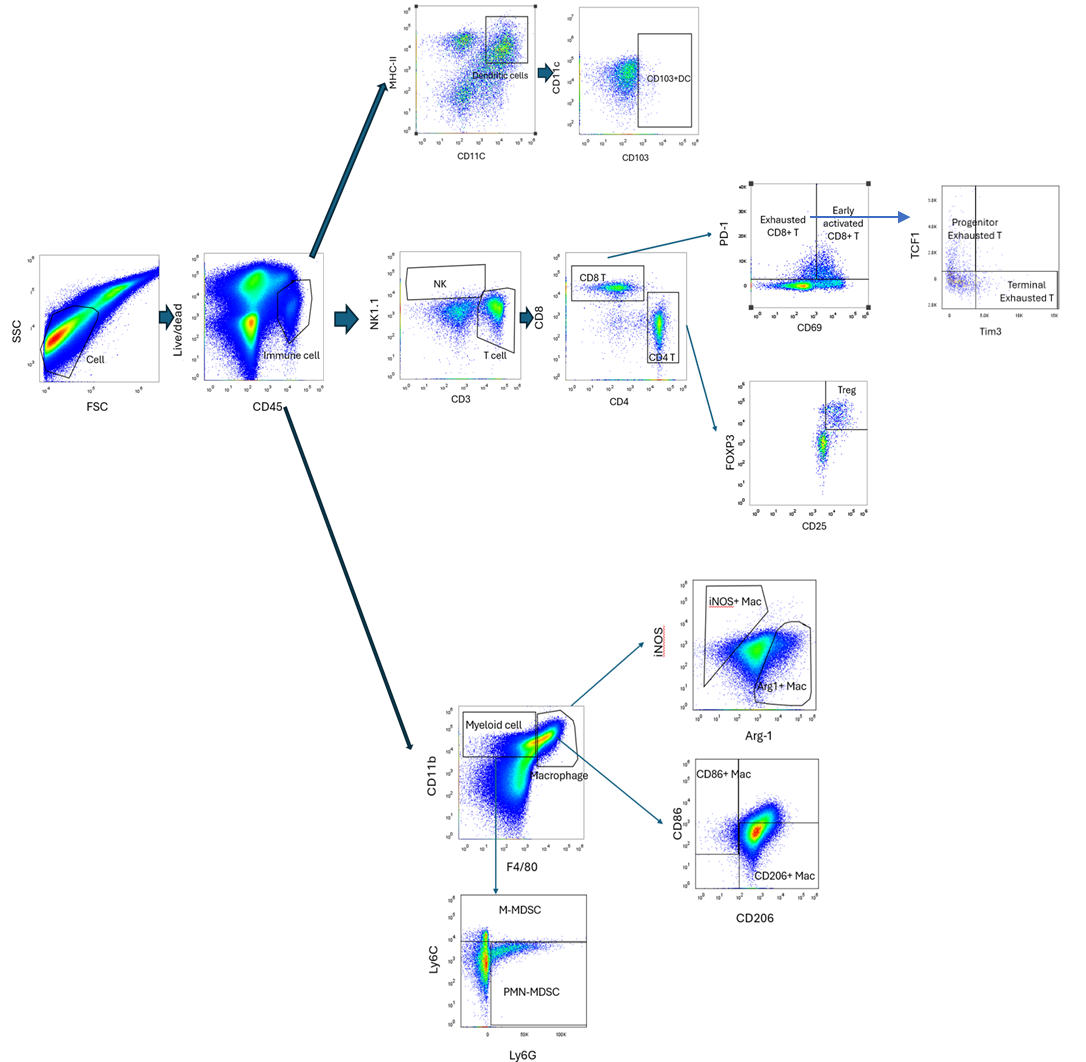

**Supplementary Fig.1. Flow cytometry gating strategy for immune cell in the tumor microenvironment.** Single-cell suspensions were gated on live CD45⁺ leukocytes to exclude debris and non-immune cells. Lymphoid and myeloid subsets were then identified. CD3⁺ T cells were divided into CD4⁺ and CD8⁺ populations; CD8⁺ T cells were analyzed for activation and exhaustion using PD-1 and CD69, they were further separated into progenitor (TCF1⁺Tim3⁻) and terminally exhausted (TCF1⁻Tim3⁺) subsets. Regulatory T cells were defined as CD4⁺CD25⁺FoxP3⁺. Dendritic cells were gated as CD11c⁺MHCII⁺, with CD103⁺ DCs as a subset. Myeloid cells (CD11b⁺) included macrophages (F4/80⁺) and MDSCs, subdivided into Ly6C⁺Ly6G⁻ (M-MDSC) and Ly6C⁻Ly6G⁺ (PMN-MDSC). Macrophage polarization was assessed by iNOS, Arg-1, CD86, and CD206 expression. Fluorescence Minus One (FMO) controls were used for each set of experiments.

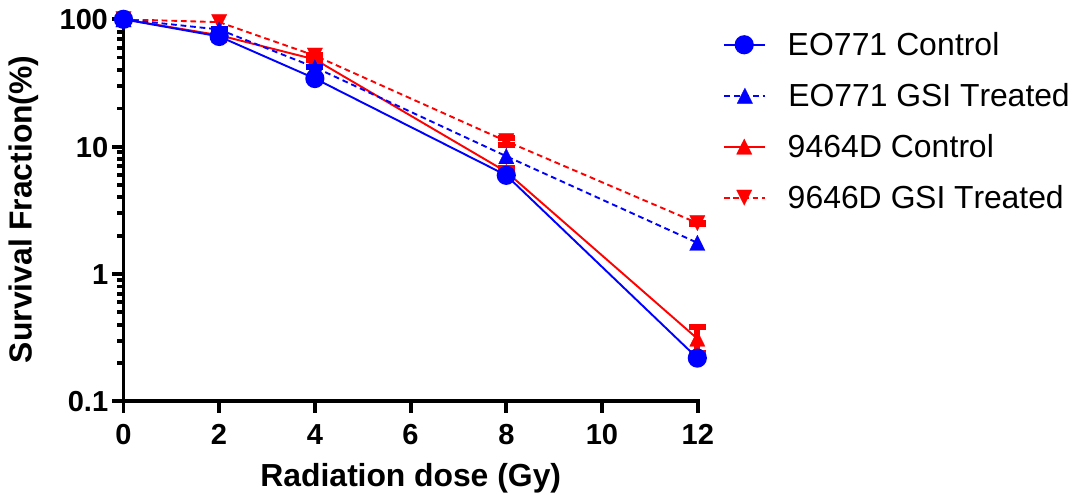

**Supplementary Fig. 2. GSI does not increase EO771 or 9464D sensitivity to RT in vitro, indicating the effect is mediated through the tumor microenvironment.** Cells were pretreated with or without 100µM GSI and radiated with 0, 2, 4, 8, and 12 Gy. No significant reduction in clonogenic survival or increased EO771 or 9464D tumor cell sensitivity to radiation was observed.
